# Estradiol Modulates Cytotoxic Responses to Topoisomerase II-Targeting Agents in Estrogen Receptor – Positive Breast Cancer Cells

**DOI:** 10.64898/2026.10.19.760233

**Authors:** Nisha Susan Thomas, Nai-Yun Sun, Mathew Cherian

## Abstract

Endocrine therapy is the primary treatment for estrogen receptor-positive (ER+) breast cancer. Paradoxically high dose estradiol can induce ER+ breast cancer cell death in an ER dependent manner. Although a well-established phenomenon, the mechanism of estradiol (E2)-induced breast cancer cell death has remained elusive. More recent evidence has revealed that E2 drives estrogen receptor alpha (ERα)-dependent transcription through TOP2 mediated DNA double-strand breaks. This raises two important questions. Firstly, is E2-dependent breast cancer cell death dependent on TOP2 mediated DNA double strand cleavage, and, secondly, does E2 signaling alter sensitivity of ER+ breast cancer cells to TOP2 poisons. Two ER+ breast cancer cell lines T47D and MCF-7 cells were treated with E2, doxorubicin (TOP2 poison) or T638 (catalytic TOP2 inhibitor), alone, and with each drug combined with E2 for 72 h. Viability was measured and normalized to untreated controls and IC_50_ values were calculated from dose-response curve. E2 enhanced doxorubicin potency in T47D and MCF7 ER+ breast cancer cell lines. E2 was toxic to T47D and MCF7 cells in a concentration dependent manner. The TOP2 catalytic inhibitor, T638, which blocks TOP2-mediated DNA cleavage reduced viability of both cell lines when used alone but decreased the cytotoxic effects of E2 about 7.6-fold, consistent with a role for TOP2-mediated DNA cleavage in the cytotoxic effects of E2. Thus, E2 sensitizes ER+ breast cancer cells to the TOP2 poison, whereas T638, the catalytic inhibitor, diminishes the cytotoxic response to E2. These opposing effects are consistent with E2 acting through TOP2-dependent DNA breaks.

## Introduction

Estrogen receptor-positive (ER+) breast cancer accounts for 70% of the newly diagnosed breast cancers (Kim & Lukong, 2025). Doxorubicin (DOX), a topoisomerase II inhibitor, is a standard chemotherapy regimen for breast cancer, including triple negative breast cancer (TNBC) and high-risk patients with ER+ breast cancer. Anthracycline-based chemotherapy lowers recurrence and mortality in early-stage breast cancer, although its benefits in ER+ breast cancer subtype are heterogenous (Early Breast Cancer Trialists’ Collaborative Group . Electronic address & Early Breast Cancer Trialists’ Collaborative, 2023). Estrogen stimulates proliferation while also modulating DNA damage response (DDR) and DNA repair pathway in ER+ tumor cells (Caldon, 2014; Ho et al., 2016). However, it’s unclear whether and how estrogen signaling enhances or limits doxorubicin mediated DNA damage response in ER+ breast cancer.

Topoisomerase 2α (TOP2A) and Topoisomerase 2β (TOP2B) creates transient DNA double stranded breaks (DSBs) to disentangle the DNA topological stress during replication, transcription and chromosome segregation (Nitiss, 2009a; Pommier et al., 2016). TOP2 poisons, doxorubicin and etoposide trap the covalent TOP2-DNA cleavage complex and prevent DNA religation. This converts the enzymes activity from creating transient intermediate double-strand breaks into a source of persistent cytotoxic DNA damage (Nitiss, 2009b). In contrast, the T60-derived compound T638, aclarubicin and suramin are catalytic inhibitors that suppress either TOP2A or both TOP2A and TOP2B function, disrupting enzyme activity by preventing TOP2 -DNA interaction, blocking ATP binding to the enzyme, or stabilizing the non-covalent TOP2-DNA complex (Matias-Barrios et al., 2021). TOP2 is a key enzyme in hormone-regulated transcription. Transient TOP2-dependent DNA breaks occur at the nuclear receptor target promoters, and PARP-1 and DNA-PK are recruited to the process. These ligand-induced breaks generate chromosomal translocations (Cowell et al., 2023; Ju & Rosenfeld, 2006; Lin et al., 2009). In breast cancer cells, estradiol (E2) triggers estrogen receptor alpha (ERα) and TOP2-dependent γH2AX foci at estrogen-responsive promoters (Williamson & Lees-Miller, 2011), and R-loop-associated, replication-dependent breaks (Stork et al., 2016). Physiological concentration of E2 (10 nM) are sufficient to induce TOP2-dependent double-strand DNA breaks in G_1_-arrested breast cancer cells and mouse mammary gland epithelial cells lacking BRCA1 (Sasanuma et al., 2018).

Early studies in T47D human epithelial breast cancer cell line showed that estrogen increases etoposide and doxorubicin induced DNA cleavage and TOP2 content. However, doxorubicin cytotoxicity was modest in these studies(Epstein & Smith, 1988). Top2A abundance is a determinant of doxorubicin response (Burgess et al., 2008) and TOP2B mediates doxorubicin cardiotoxicity (Yeh & Chang, 2025), indicating that both TOP2A and TOP2B together shapes the therapeutic efficacy and tolerability. We therefore determined whether estrogen signaling differentially influences ER+ breast cancer cells treated with TOP2 poison and a catalytic inhibitor.

## Materials and Methods

### Cell lines and culture

Human ER+ breast cancer cell lines MCF-7 (HTB-22) and T-47D (HTB-133) were purchased from ATCC and were maintained according to ATCC recommendation. MCF-7 cells were cultured in Dulbecco’s modified eagle medium (DMEM) supplemented with 0.01 mg/mL human recombinant insulin and 10% fetal bovine serum (FBS). T-47D cells were cultured in RPMI-1640 medium supplemented 10% FBS. Cells were grown at 37ºC in a humidified atmosphere of 5% CO_2_ in air. Cells were passaged at 70% confluence, land the media was renewed 2 days once. MCF-7 cells were seeded at 5,000 cells per well in 100 µL media in 96 well plates and T-47D cells were seeded at 2,500 cells per well in 100 µL medium. Before hormone treatments, MCF-7 and T-47D cells were washed with 1x phosphate-buffered saline (PBS) and hormone deprived in phenol red-free Dulbecco’s modified eagle medium (DMEM) or RPMI-1640 containing 0.5% charcoal-stripped fetal bovine serum for 16h to reduce steroid receptor activation. All the treatments were performed for 72h in this media.

### Reagents and treatments

17β-Estradiol ((E2); Selleck chemical LLC# S1709), doxorubicin hydrochloride (Tocris Bioscience# 2252) and T638 (MedChem Express# HY-164143) were dissolved in DMSO and diluted in medium. The final vehicle concentration was 0.1%.

#### Doxorubicin Dose-Response Experiment

Doxorubicin was serially diluted in phenol free 0.5% charcoal stripped FBS containing DMEM or RPMI-1640 media. MCF-7 cells were treated with doxorubicin at 5, 2.5, 1.25, 0.625, 0.312, 0.156, 0.078, 0.039 or 0.019.5 µM and T47D cells at 1, 0.5, 0.25, 0.125, 0.0625, 0.03125, 0.0156, 0.0078 or 0.0039 µM.

#### Estradiol Dose-Response Experiments

For E2 alone experiments, cells were serum starved for 16h in phenol free media containing 0.5% charcoal stripped FBS followed by E2 treatment. MCF-7 cells were treated with 10, 1, 0.1, 0.01 or 0.001 µM and T47D cells at 0.1, 0.01 or 0.001 µM.

#### T638 Dose-Response Experiment

Both MCF-7 and T47D cells were treated with 1, 0.5 or 0.1 µM of T638 in in phenol free media containing 0.5% charcoal stripped FBS.

#### Combination experiments

For the drug combinations, the same cell lines were treated in the same media with combinations of the drugs used above in the same concentration range.

### Cell Viability Assay

Cell viability was determined using PrestoBlue™ HS cell viability reagent (Invitrogen, Thermo Fisher Scientific #P50201). At the end of treatment period, 10 ul of reagent was added to each well. Plates were protected from light and incubated at 37ºC in a humidified 5% CO2 incubator for 30 min. Fluorescence intensity was measured on SpectraMax iD5e reader at excitation wavelength 560 nm and emission wavelength 600 nm. Fluorescence values were background corrected using cell free medium blanks and normalized to vehicle-treated controls and expressed as percentage of vehicle-treated control viability.

### Data analysis

Dose-response data were fitted with a four-parameter nonlinear regression of inhibitor vs. response analyzed in GraphPad Prism 10 to obtain the IC_50_ value. Viability was normalized to vehicle-treated control and presented as a percentage. The 95% confidence intervals were calculated by the profile likelihood method. Fold-shift was calculated as IC_50_ (single agent) ÷ IC_50_ (combination) or the reverse for rescue. Data are mean ± SEM of 3 independent experiments with 3 replicates.

## Results

### Estrogen and doxorubicin reduce ER+ breast cancer cell viability with varying sensitivities

Increasing concentrations of E2 were associated with decreased cell viability of both T-47D and MCF-7 cell lines. The cell viability of T-47D was reduced with an IC_50_ of 0.0275 µM (Fig. 1A) and that of MCF7 cells 0.4895 µM (Fig. 1D), showing that T-47D cells were about 18-fold more sensitive to E2 as compared to MCF-7 cells. Treatment with doxorubicin alone also caused concentration-dependent viability loss, with an IC_50_ of 0.234 µM in T47D (Fig.1B) and 2.22 µM in MCF-7 cells (Fig. 1E). T47D cells were about 9.5-fold more sensitive to doxorubicin.

**Figure 1.**
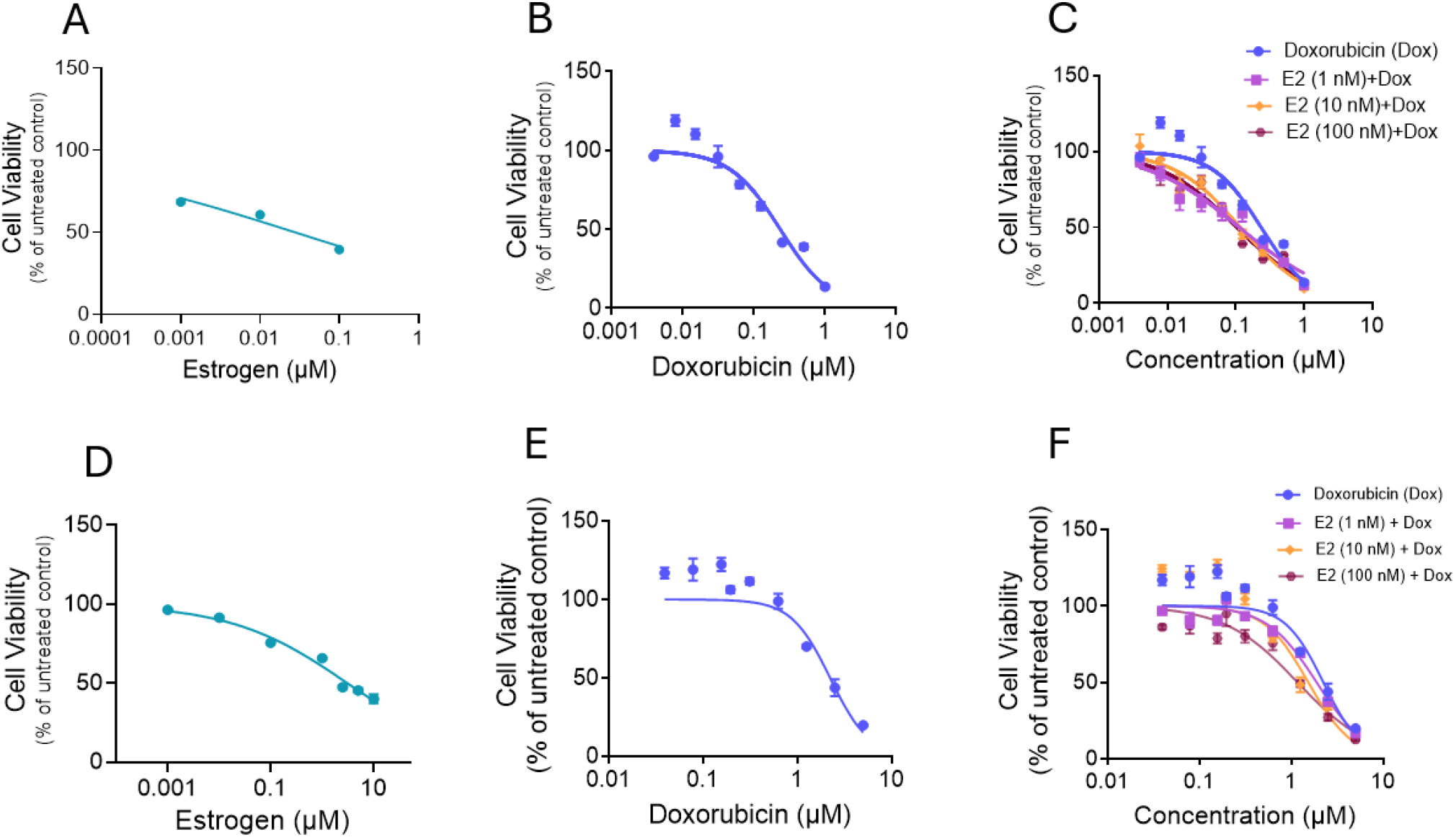
Estradiol enhances the cytotoxic effect of doxorubicin in ER+ breast cancer cells. (A, D) Dose-response curves for the T47D and MCF-7 cells treated with E2 alone (IC50; 0.0275 µM and 0.4895 μM, respectively). (B,E) Dose Response curve for T47D and MCF-7 cells treated with Doxorubicin alone (IC50; 0.234 µM and 2.22 µM, respectively). (C,F) Graph representing treatment with doxorubicin combined with E2 at 1,10 or 100 nM. Cell viability is expressed as percentage of the untreated control. Nonlinear regression was used to estimate half maximal inhibitory concentrations (IC50). Data are mean ± SEM of 3 independent experiments, treated for 72 h.

### Estradiol enhances the cytotoxicity of doxorubicin

E2 at doses 0.1, 0.01 or 0.001 µM consistently enhanced doxorubicin potency on ER+ breast cancer cell viability, as shown by leftwards shift of the dose-response curves in both T47D and MCF7 ER+ breast cancer cell lines. In T-47D cells (Fig. 1C), E2 lowered the doxorubicin IC_50_ 2.1 (1nM E2), 2.0 (10nM E2) and 2.3-fold (100nM E2. Hence, the effect was maximal at 1 nM and did not increase significantly at 10 or 100 nM. In MCF-7 cells (Fig. 1F), doxorubicin sensitization depends on the E2 concentration. The IC_50_ of doxorubicin decreased from 2.22 µM (95% CI 1.757–2.850) to 1.86 (95% CI 1.672-2.116), 1.52 (95% CI 1.129-2.109) and 1.116 µM (95% CI 0.9430-1.446) with 0.001, 0.01 or 0.1 µM E2, a 1.2-,1.5- and 1.9-fold increase in doxorubicin potency. Hence, the increase in doxorubicin potency in MCF-7 cells increased with increasing E2 concentrations.

### TOP2 catalytic inhibitor T638 alone reduces the cell viability

Treatment of T638 alone in T-47D and MCF-7 (Fig. 2B, E) cells decreased viability with an IC_50_ of 0.598 µM in T-47D cells and 0.129 µM in MCF-7 cells, showing that MCF-7 cells were 4.6-fold more sensitive.

**Figure 2.**
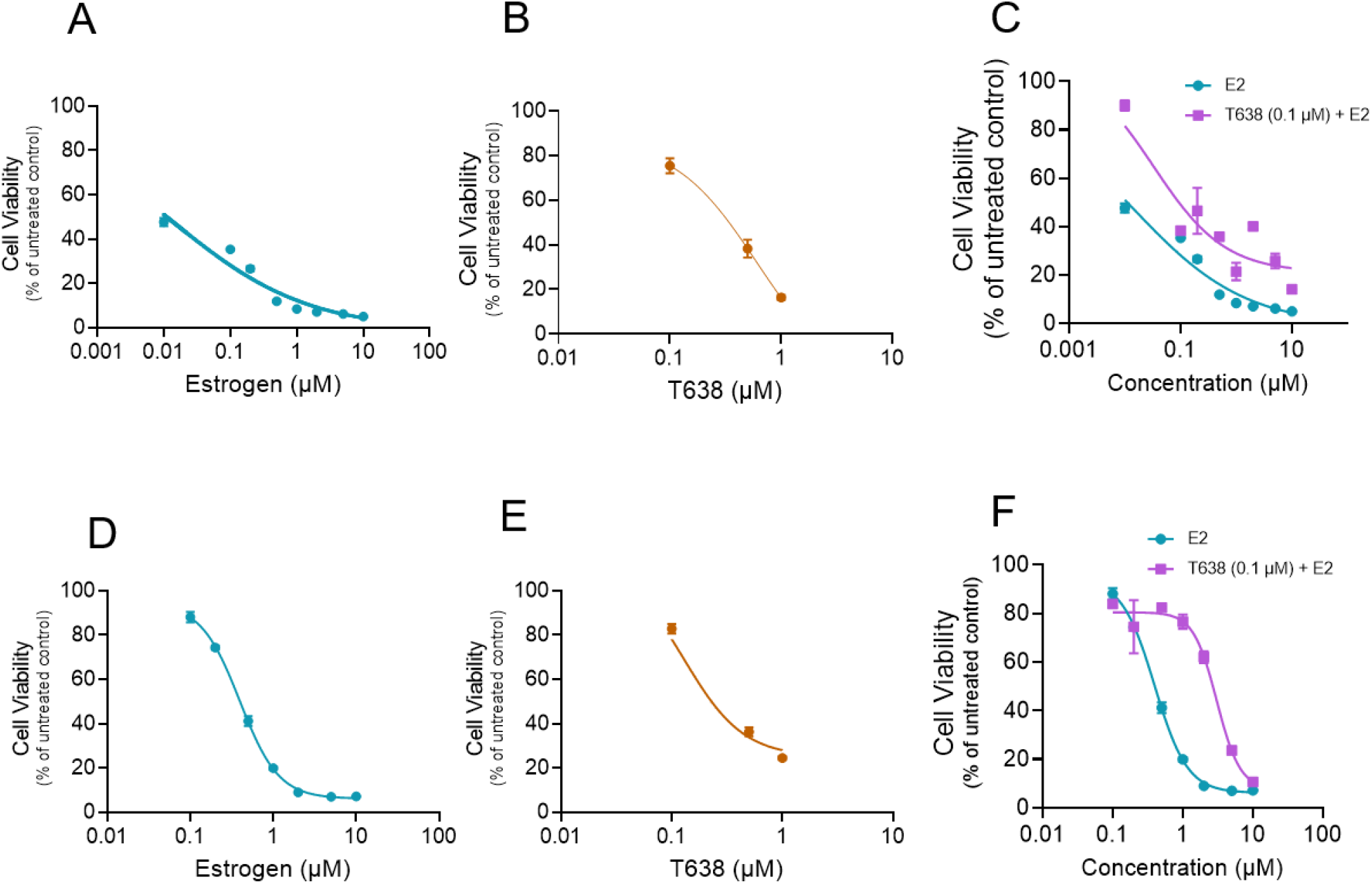
Catalytic TOP2 inhibitor T638 attenuates E2-induced cell viability loss in ER+ breast cancer cells. (A,D) Dose response curve for E2 alone treated T47D and MCF-7 (IC50; 0.01407 µM and 0.3962 μM, respectively). (B.E) Dose-response curve for T638 alone treated T47D and MCF-7 cells (IC50; 0.5983 µM and 0.1292 µM, respectively). (C.F) E2 and T638 combination effect in T47D and MC-7 cells showing T638 increases the cell viability in both the cell when combined with E2 as compared to E2 alone treated cells. Viability is expressed as a percentage of untreated control. Data are mean ± SEM of 3 independent experiments, treated for 72 h.

### T638 attenuates estradiol induced loss of cell viability

Treatment with E2 alone (Fig. 2A) resulted in loss of viability with IC_50_ of 0.01407 µM in T47D cells and 0.3962 µM in MCF-7 cells (Fig. 2D). Whereas concurrent treatment with E2 and T638 at 0.1 µM shifted the E2 response curve to the right, showing more cells remained viable at equivalent E2 concentrations (Fig. 2C, F). In T-47D cells the IC_50_ value of E2 increased from 0.0141 µM (95% CI 0.00715-0.1874) to 0.032 µM (95% CI 0.01742-0.1043) showing a 2.3-fold change. In MCF-7 cells the IC_50_ value increased from 0.3962 µM (95% CI 0.3068-0.4349) to 3.050 µM (95%CI 1.949).

## Discussion

In this study, estradiol (E2) and TOP2 drugs showed opposing effects in the ER+ breast cancer cells. E2 increased the cytotoxicity of TOP2 poison doxorubicin in both the lines, whereas the T638, the TOP2 catalytic inhibitor reduced E2-induced viability loss. These results suggest the cytotoxic effect of E2 on ER+ breast cancer cells is TOP2 dependent. This effect could be due to the role of TOP2 in hormone regulated transcription. Previous studies have shown that estrogen mediated estrogen receptor signaling recruits TOP2and creates transient breaks at ER target genes (Williamson & Lees-Miller, 2011). Hormone induced TOP2mediated DNA double strand breaks have been proposed as a therapeutic target in prostate cancer (Haffner et al., 2010). These studies demonstrate that E2 increases the number of TOP2 cleavage events in the cells. Doxorubicin, a TOP2 poison, can stabilize these otherwise transient lesions (Pommier et al., 2010; Tewey et al., 1984), which is consistent with our study findings showing a shift in doxorubicin sensitivity when combined with E2. On the other hand, TOP2 catalytic inhibitors, which block the capacity of TOP2 to cleave DNA (Matias-Barrios et al., 2021; Matias-Barrios et al., 2020). Our findings show that the TOP2 catalytic inhibitor T638 decreases the cytotoxic effects of E2 in T47D and MCF7 ER+ breast cancer cells.

Our results build on earlier work in T-47D cells, in which estrogen increases TOP2 poison-induced DNA cleavage, and novobiocin, a catalytic TOP2 inhibitor, antagonized the effect on the DNA cleavage (Epstein & Smith, 1988; Epstein et al., 1989). That study found only minimal enhancement of doxorubicin cytotoxicity, whereas we observed an approximately 2-fold IC_50_ shift after 72 h of treatment. The two cells lines responded differently to concurrent E2 treatment in terms of incremental sensitivity to higher E2 concentrations. T47D cells were more sensitive to E2 and doxorubicin, achieving maximal sensitivity at 1 nM E2. MCF-7 were less sensitive but showed dose-dependent sensitization across the E2 range. In contrast, MCF-7 cells were more sensitive to T638 and showed a larger T638 effect on E2 response, about 7.6-fold versus 2.3-fold in T47D cells. TOP2A abundance is a determinant of doxorubicin response (Burgess et al., 2008) and correlates with ER+ tumor proliferation (Rody et al., 2009; Tokiniwa et al., 2012). Differences in the ERα expression, proliferation rate or TOP2 isoform expression levels may explain the differences observed between the two ER+ breast cancer cell lines.

Clinically anthracycline benefit in ER+ cancer is heterogeneous and HER2 and TOP2A are only modest predictors (Di Leo et al., 2011; Early Breast Cancer Trialists’ Collaborative Group. Electronic address & Early Breast Cancer Trialists’ Collaborative, 2023) Based on our findings, concurrent E2 treatment could sensitize these cancers to anthracyclines.

## Conclusion

E2 enhances the sensitivity of ER+ breast cancer cells to the TOP2 poison doxorubicin, while the catalytic TOP2 inhibitor T638 attenuates E2-induced cytotoxicity. These findings are consistent with the role for TOP2-dependent DNA break formation in the cytotoxic effects of E2 and for synergy between E2 and TOP2 poisons in ER+ breast cancer cell lines. Further mechanistic and *in vivo* studies using knockout models are required to determine whether E2 could be used to increase treatment response in ER+ breast cancer.

## References

Burgess, D. J., Doles, J., Zender, L., Xue, W., Ma, B., McCombie, W. R., Hannon, G. J., Lowe, S. W., & Hemann, M. T. (2008). Topoisomerase levels determine chemotherapy response in vitro and in vivo. Proc Natl Acad Sci U S A, 105(26), 9053–9058. 10.1073/pnas.0803513105

Caldon, C. E. (2014). Estrogen signaling and the DNA damage response in hormone dependent breast cancers. Front Oncol, 4, 106. 10.3389/fonc.2014.00106

Cowell, I. G., Casement, J. W., & Austin, C. A. (2023). To Break or Not to Break: The Role of TOP2B in Transcription. Int J Mol Sci, 24(19). 10.3390/ijms241914806

Di Leo, A., Desmedt, C., Bartlett, J. M., Piette, F., Ejlertsen, B., Pritchard, K. I., Larsimont, D., Poole, C., Isola, J., Earl, H., Mouridsen, H., O’Malley, F. P., Cardoso, F., Tanner, M., Munro, A., Twelves, C. J., Sotiriou, C., Shepherd, L., Cameron, D.,…Group, H. T. A. M.-a. S. (2011). HER2 and TOP2A as predictive markers for anthracycline-containing chemotherapy regimens as adjuvant treatment of breast cancer: a meta-analysis of individual patient data. Lancet Oncol, 12(12), 1134–1142. 10.1016/S1470-2045(11)70231-5

Early Breast Cancer Trialists’ Collaborative Group . Electronic address, b. o. c. o. a. u., & Early Breast Cancer Trialists’ Collaborative, G. (2023). Anthracycline-containing and taxane-containing chemotherapy for early-stage operable breast cancer: a patient-level meta-analysis of 100 000 women from 86 randomised trials. Lancet, 401(10384), 1277–1292. 10.1016/S0140-6736(23)00285-4

Epstein, R. J., & Smith, P. J. (1988). Estrogen-induced potentiation of DNA damage and cytotoxicity in human breast cancer cells treated with topoisomerase II-interactive antitumor drugs. Cancer Res, 48(2), 297–303.

Epstein, R. J., Smith, P. J., Watson, J. V., Waters, C., & Bleehen, N. M. (1989). Oestrogen potentiates topoisomerase-II-mediated cytotoxicity in an activated subpopulation of human breast cancer cells: implications for cytotoxic drug resistance in solid tumours. Int J Cancer, 44(3), 501–505. 10.1002/ijc.2910440321

Haffner, M. C., Aryee, M. J., Toubaji, A., Esopi, D. M., Albadine, R., Gurel, B., Isaacs, W. B., Bova, G. S., Liu, W., Xu, J., Meeker, A. K., Netto, G., De Marzo, A. M., Nelson, W. G., & Yegnasubramanian, S. (2010). Androgen-induced TOP2B-mediated double-strand breaks and prostate cancer gene rearrangements. Nat Genet, 42(8), 668–675. 10.1038/ng.613

Ho, J. Y., Chang, F. W., Huang, F. S., Liu, J. M., Liu, Y. P., Chen, S. P., Liu, Y. L., Cheng, K. C., Yu, C. P., & Hsu, R. J. (2016). Estrogen Enhances the Cell Viability and Motility of Breast Cancer Cells through the ERalpha-DeltaNp63-Integrin beta4 Signaling Pathway. PLoS One, 11(2), e0148301. 10.1371/journal.pone.0148301

Ju, B. G., & Rosenfeld, M. G. (2006). A breaking strategy for topoisomerase IIbeta/PARP-1-dependent regulated transcription. Cell Cycle, 5(22), 2557–2560. 10.4161/cc.5.22.3497

Kim, N., & Lukong, K. E. (2025). Treating ER-positive breast cancer: a review of the current FDA-approved SERMs and SERDs and their mechanisms of action. Oncol Rev, 19, 1564642. 10.3389/or.2025.1564642

Lin, C., Yang, L., Tanasa, B., Hutt, K., Ju, B. G., Ohgi, K., Zhang, J., Rose, D. W., Fu, X. D., Glass, C. K., & Rosenfeld, M. G. (2009). Nuclear receptor-induced chromosomal proximity and DNA breaks underlie specific translocations in cancer. Cell, 139(6), 1069–1083. 10.1016/j.cell.2009.11.030

Matias-Barrios, V. M., Radaeva, M., Ho, C. H., Lee, J., Adomat, H., Lallous, N., Cherkasov, A., & Dong, X. (2021). Optimization of New Catalytic Topoisomerase II Inhibitors as an Anti-Cancer Therapy. Cancers (Basel), 13(15). 10.3390/cancers13153675

Matias-Barrios, V. M., Radaeva, M., Song, Y., Alperstein, Z., Lee, A. R., Schmitt, V., Lee, J., Ban, F., Xie, N., Qi, J., Lallous, N., Gleave, M. E., Cherkasov, A., & Dong, X. (2020). Discovery of New Catalytic Topoisomerase II Inhibitors for Anticancer Therapeutics. Front Oncol, 10, 633142. 10.3389/fonc.2020.633142

Nitiss, J. L. (2009a). DNA topoisomerase II and its growing repertoire of biological functions. Nat Rev Cancer, 9(5), 327–337. 10.1038/nrc2608

Nitiss, J. L. (2009b). Targeting DNA topoisomerase II in cancer chemotherapy. Nat Rev Cancer, 9(5), 338–350. 10.1038/nrc2607

Pommier, Y., Leo, E., Zhang, H., & Marchand, C. (2010). DNA topoisomerases and their poisoning by anticancer and antibacterial drugs. Chem Biol, 17(5), 421–433. 10.1016/j.chembiol.2010.04.012

Pommier, Y., Sun, Y., Huang, S. N., & Nitiss, J. L. (2016). Roles of eukaryotic topoisomerases in transcription, replication and genomic stability. Nat Rev Mol Cell Biol, 17(11), 703–721. 10.1038/nrm.2016.111

Rody, A., Karn, T., Ruckhaberle, E., Muller, V., Gehrmann, M., Solbach, C., Ahr, A., Gatje, R., Holtrich, U., & Kaufmann, M. (2009). Gene expression of topoisomerase II alpha (TOP2A) by microarray analysis is highly prognostic in estrogen receptor (ER) positive breast cancer. Breast Cancer Res Treat, 113(3), 457–466. 10.1007/s10549-008-9964-x

Sasanuma, H., Tsuda, M., Morimoto, S., Saha, L. K., Rahman, M. M., Kiyooka, Y., Fujiike, H., Cherniack, A. D., Itou, J., Callen Moreu, E., Toi, M., Nakada, S., Tanaka, H., Tsutsui, K., Yamada, S., Nussenzweig, A., & Takeda, S. (2018). BRCA1 ensures genome integrity by eliminating estrogen-induced pathological topoisomerase II-DNA complexes. Proc Natl Acad Sci U S A, 115(45), E10642–E10651. 10.1073/pnas.1803177115

Stork, C. T., Bocek, M., Crossley, M. P., Sollier, J., Sanz, L. A., Chedin, F., Swigut, T., & Cimprich, K. A. (2016). Co-transcriptional R-loops are the main cause of estrogen-induced DNA damage. Elife, 5. 10.7554/eLife.17548

Tewey, K. M., Rowe, T. C., Yang, L., Halligan, B. D., & Liu, L. F. (1984). Adriamycin-induced DNA damage mediated by mammalian DNA topoisomerase II. Science, 226(4673), 466–468. 10.1126/science.6093249

Tokiniwa, H., Horiguchi, J., Takata, D., Kikuchi, M., Rokutanda, N., Nagaoka, R., Sato, A., Odawara, H., Tozuka, K., Oyama, T., & Takeyoshi, I. (2012). Topoisomerase II alpha expression and the Ki-67 labeling index correlate with prognostic factors in estrogen receptor-positive and human epidermal growth factor type-2-negative breast cancer. Breast Cancer, 19(4), 309–314. 10.1007/s12282-011-0291-4

Williamson, L. M., & Lees-Miller, S. P. (2011). Estrogen receptor alpha-mediated transcription induces cell cycle-dependent DNA double-strand breaks. Carcinogenesis, 32(3), 279–285. 10.1093/carcin/bgq255

Yeh, E. T. H., & Chang, H. M. (2025). Anthracycline-Induced Cardiotoxicity: Mechanism and Prevention. Trans Am Clin Climatol Assoc, 135, 184–195.

